# α5-GABA-A Receptor Positive Allosteric Modulation prevents neuronal atrophy and cognitive decline independently of p-Tau accumulation in the PS19 mouse model

**DOI:** 10.1101/2024.09.07.611810

**Authors:** Ravinder N. Dharavath, Ashley M. Bernardo, Cassandra Marceau-Linhares, Michael Marcotte, Carla Mezo-Gonzalez, Kayla Wong, Celeste Pina-Leblanc, Adrien Bouchet, Dishary Sharmin, Kamal P. Pandey, James M. Cook, Thomas D. Prevot, Etienne Sibille

**Affiliations:** Campbell Family Mental Health Research Institute of CAMH, 250 college street, Toronto, ON, M5T 1R8 Canada; Department of Chemistry and Biochemistry, University of Wisconsin–Milwaukee, 3210 N Cramer Street, 53211, WI, USA; Department of Psychiatry, University of Toronto, 250 college street, Toronto, ON, M5T 1R8 Canada; Department of Pharmacology and Toxicology, University of Toronto, Medical Sciences Building, 1 King’s College Circle Room 4207, Toronto, ON, M5S 1A8, Canada

**Author notes:** Corresponding Authors: Etienne Sibille, Ph.D, CAMH, 250 College Street, room 134, Toronto, ON M5T 1R8, Canada; Thomas D. Prevot, Ph.D., CAMH, 250 College Street, room 131, Toronto, ON M5T 1R8, Canada. These authors equally contributed to the study (Ravinder N. Dharavath, Ashley M. Bernardo, Cassandra Marceau-Linhares).

## Abstract

**Background:** Dysregulated Tau phosphorylation (p-Tau) is a hallmark of neurodegenerative disorders such as Alzheimer’s disease (AD) or frontotemporal dementia (FTD), resulting in neurofibrillary tangle accumulation, neuronal atrophy and cognitive impairment. Impaired somatostatin (SST) expression and reduced SST-expressing GABAergic neurons significantly contributes to AD-related pathophysiology and correlates with cognitive impairment. SST+ interneurons inhibit the dendrites of excitatory neurons in cortical layers and hippocampus, primarily through α5-GABA-A receptors, regulating cognitive function. Leveraging a newly developed small molecule that targets the α5-GABA-A receptors via positive allosteric modulation (α5-PAM), we assessed its effects on p-Tau-related neuronal morphology, cognitive deficits and protein expression.

**Methods:** In the PS19 transgenic mouse model, we administered the α5-PAM, GL-II-73, either acutely or chronically at 3 and 6 months, corresponding to early and advanced stage of p-Tau accumulation. Golgi staining analyzed dendritic morphology and spine density in mice chronically exposed to α5-PAM. Western blotting was used to quantify p-tau and Tau expression. Spatial working memory was assessed using the Y-maze.

**Results:** Chronic treatment at 3 and 6 months mitigated p-Tau-induced loss of spine density and reduced dendritic length. α5-PAM treatment did not affect p-tau levels. α5-PAM effectively reversed spatial working memory deficits induced by p-tau accumulation both acutely and chronically.

**Conclusions:** α5-GABA-A receptor positive allosteric modulation displayed neurotrophic (spine and dendritic pathology) and procognitive (working memory) effects in the PS19 model, independently of p-Tau burden. This suggests a novel therapeutic strategy for p-Tau-related pathologies with both symptomatic and disease-modifying potential.

## INTRODUCTION

Tauopathies are a group of progressive neurodegenerative disorders marked by the pathological accumulation of hyperphosphorylated tau protein (p-Tau), which aggregates into neurofibrillary tangles (NFTs) (*1*), and lead to neuronal dysfunctions, cognitive deficits, and brain atrophy (*2*). Alzheimer’s disease (AD) and frontotemporal dementia (FTD) are amongst the most common tauopathies and share these pathological hallmarks (*3, 4*). The accumulation of p-Tau correlates with the progression of neuronal atrophy and cognitive deficits progression, contributing substantially to disease burden and progression (*5*).

Despite clear understanding of the role of p-Tau aggregates and NFTs in the pathophysiology of AD and other tauopathy-based disorders, pharmacotherapies targeting Tau have had limited success and interest, likely due to the failures in the AD field, when targeting Tau or amyloid β (*6–8*). However, the recent approval of antibody-based therapies targeting the soluble and insoluble amyloid protein (*9*), raised interest in revisiting both amyloid β-targeting and Tau-targeting interventions. Therapies targeting the Tau-pathology include Zagotenemab (a humanized anti-Tau antibody(*10*)), LMTX (TauRx, an aggregation inhibitor (*11*)), TPI-287 (a microtubule stabilizer, (*12*)) and others (*13*), but so far none have demonstrated efficacy at resolving neuronal atrophy or cognitive decline in patients.

Although p-Tau accumulation is a major contributor to AD and FTD pathophysiologies, neurodegenerative disorders are complex and multifactorial diseases that may require a polypharmacopeia approach to reduce the burden of the disease and improve patients’ quality of life (*14*). Aβ- and Tau-targeting therapies can help with clearing or limiting accumulation of plaques and tangles, but no studies demonstrated better neuronal function after Aβ clearance or tangle accumulation inhibition following these therapies. Neuronal hyperexcitability is consistently reported in the hippocampus (*15*) and prefrontal cortex (*16*) (brain regions highly involved in cognitive functions (*17*)) of AD patients, and proposed to be associated with Tau-accumulation (*18*). This hyperexcitability is thought to be due in part to a loss of GABAergic inhibition (*19*). Evidence supports the contribution of GABAergic inhibition in memory (*20, 21*) and demonstrated that altered GABAergic signaling is linked to poorer cognitive functions in elderly (*22*), in AD (*23*) and in FTD (*24*) patients. Therefore, targeting the GABAergic system is emerging as a valid approach for reducing biological changes in AD pathology, potentially providing disease-modifying effects and symptom relief.

Recently, the α5-GABAA receptors (α5-GABAA-R) have been proposed as relevant targets for cognitive regulation (*25, 26*), aligned with their discrete distribution in the hippocampus and prefrontal cortex. α5-GABAA-Rs are localized synaptically and extrasynaptically in distal dendrites of pyramidal neurons. In patients with AD, α5-GABAA-R binding was reduced by∼30% reduction in the hippocampus (*27*), the associated gene (*GABRA5*) was transcriptional downregulated in middle temporal gyrus (*28*). An especially promising treatment involves the exploration of compounds selectively modulating α5-GABAA-R (*26*). Previous studies in our group showed that treatment with GL-II-73, a novel α5-GABAA-R positive allosteric modulator (α5-PAM), reverses cognitive deficits in animal models of stress (*29, 30*), aging (*31*), and amyloid deposition (*32*). In addition, we showed that chronic GL-II-73 treatment reversed the loss of dendritic lengths, spine counts and spine density in the prefrontal cortex (PFC) and hippocampus of animals in the same three models of these conditions (*30–32*).

In the present study, we hypothesize that chronic α5-PAM treatment prevents and/or reverses neuronal deficits during the early, and potentially later stage, of p-Tau accumulation, and that these effects would be associated with improved cognitive functions. To test this hypothesis, we investigated the potential of GL-II-73 to mitigate cognitive deficits and neuronal loss in the PS19 mouse model, characterized by the progressive accumulation of p-Tau. The PS19 model expresses the P301S mutant form of human microtubule-associated protein Tau (MAPT), under the control of the mouse prion protein promoter (Prnp) (*33*). These mice exhibit increased accumulation of p-Tau, leading to cognitive decline and neuronal damage (*34*). We conducted assessments on both male and female mice at 3 and 6 months of age, representing early and later stages of Tau tangle deposition.

## METHODS

### Ethical Statement

All animal procedures adhered to the Ontario Animals for Research Act (RSO 1990, Chapter A.22) and the Canadian Council on Animal Care (CCAC). The study was approved by the Institutes Animal Care Committees (Protocol #846).

### Animals

PS19 breeders, expressing the P301S mutant form of human microtubule-associated protein Tau (MAPT), were obtained from Jackson Laboratories (Cat #008169-JAX) on a C57BL/6xC3H background and paired with WT breeders to generate experimental groups’ parents. Four cohorts of PS19 heterozygous (HET) and WT littermates were produced: Acute studies at 3 months (N=8 per group, 3 groups 50% male, **Fig. 1A**) and 6 months (N=8-9 per group, 3 groups, ∼50% female, **Fig. 1B**); Chronic studies at 2-3 months (N=12-13 per group, 3 groups, 50% female, **Fig. 1C**) and 5-6 months (N=12-13 per group, 3 groups, 50% female, **Fig. 1D**). Animals were group-housed until the study commenced, then transitioned to single housing to facilitate drug administration and monitoring. They had ad libitum access to food and water and were maintained on a 12-hour light/dark cycle. Animal handling followed the technique outlined in Marcotte et al (*35*) to minimize reactivity to the experimenter.

**Figure 1.**
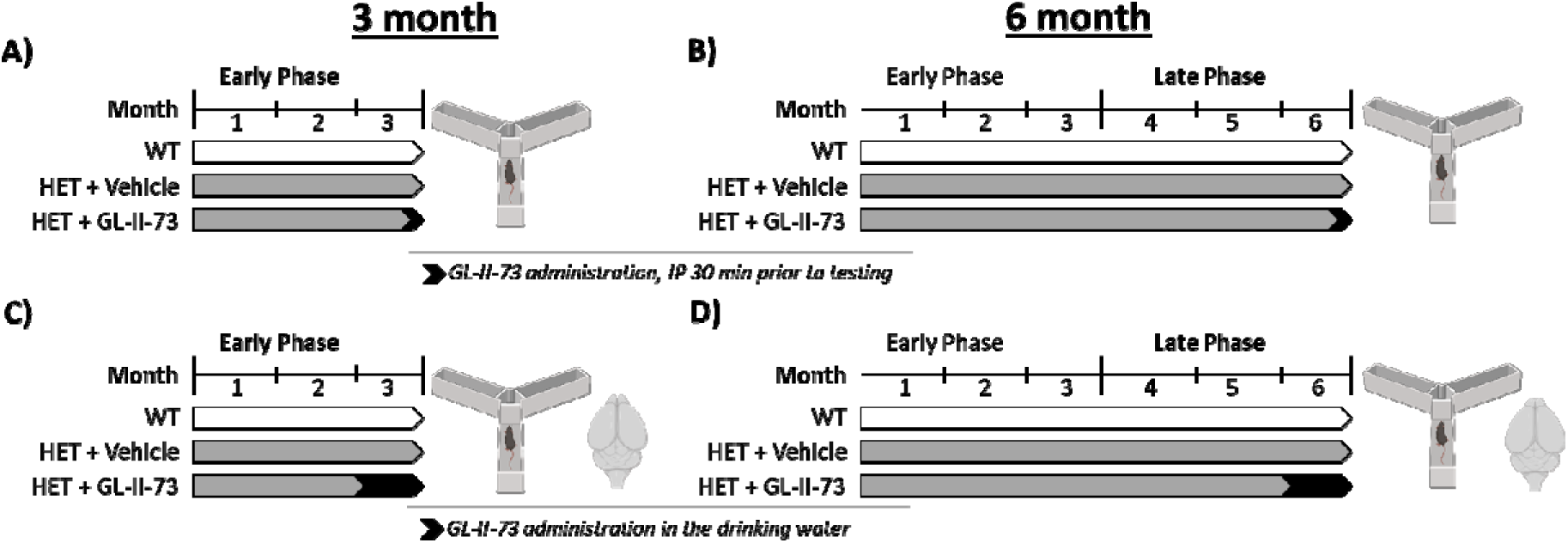
Study Design. Heterozygote PS19 mice (HET) and their wildtype (WT) littermates were included in these studies, assessing the effect of GL-II-73 on working memory and neuronal morphology. Acute treatment (**A-B**) was provided to the animals via intraperitoneal administration at 5mg/kg or 10mg/kg at 3 (**A**) or 6 (**B**) months of age. Chronic administration of GL-II-73 was also investigated at 3 (**C**) and 6 (**D**) months, with 4 weeks of administration in the drinking water at 30 mg/kg. One week after completion of the behavioral screen, mice were euthanized by rapid cervical dislocation. Brains from 4 mice per group were collected for Golgi staining, while the remaining 8 per group were collected for Western Blot analyses.

### Drug Preparation and Administration

GL-II-73, an α5-GABAA receptor (α5-GABAA-R) positive allosteric modulator (PAM) (*29*), was synthesized by Dr. Cook’s group (University of Wisconsin-Milwaukee). For acute administration via intraperitoneal (IP) injection, GL-II-73 was dissolved in a vehicle solution (85% distilled H2O, 14% propylene glycol (Sigma Aldrich), 1% Tween 80 (Sigma Aldrich)). Acute studies at 3 months and 6 months utilized GL-II-73 at 5mg/kg and 10mg/kg in the Y maze via IP injection, administered 30 minutes before testing. For chronic studies, GL-II-73 was dissolved in tap water and orally administered to mice at 30mg/kg in the drinking water, based on daily water intake calculations as in (*36*). Drug solution dissolved in drinking water was prepared fresh every other day and was provided ad libitum for 4 weeks.

### Y-Maze

The Y maze, adapted from Faucher et al. (*37*), uses spontaneous alternation to assess spatial working memory. It consists of three arms—two “goal” arms and one “starting” arm—separated by 120°, with sliding doors for the goal arms and a start box at the end of the starting arm. Mice undergo a 2-day habituation phase, exploring the maze for 10 minutes each day. On the third day, the training phase begins. Mice are placed in the start box, and after a 30-second interval, the sliding door opens, allowing them to choose a goal arm. After selection, the goal arm’s door closes, and the mouse stays there for 30 seconds before being returned to the start box. This is repeated for seven trials. On the fourth day, the procedure is repeated with a 60-second interval between trials, except for the eighth trial, where a 10-second interval tests motivation to alternate. Mice that do not alternate in the eighth trial are excluded to prevent motivational bias.

Mean % alternation is calculated as [(Number of alternations/number of trials) * 100]. The maze is cleaned with 70% ethanol between trials to remove olfactory cues.

### Golgi Staining and Analysis

After euthanasia and brain extraction, brains from six mice per group (three males, three females) were placed in Golgi Solution for two weeks, selected pseudo-randomly based on Y maze performance. They were then coded for blinded analysis and sent to NeuroDigitech (San Diego, CA, USA). Remaining brains were dissected for PFC and hippocampus extraction for western blot analysis.

For Golgi-stained brains, 100µm coronal slices were made using a cryostat and mounted on glass slides. Basal and apical dendrites of pyramidal neurons in layers II/III of the PFC and CA1 were identified as regions of interest (ROIs). Dendritic features were quantified using NeuroLucida v10 on a Zeiss Axioplan 2 microscope with an Optronics MicroFire CCD camera. Motorized X, Y, and Z focus enabled high-resolution imaging. Low magnification identified ROIs, while higher magnifications (40x-60x) were used for detailed analysis. Immersion oil and a Zeiss 100x lens facilitated 3D imaging and spine counting. Neurons (n=6 per animal) were selected based on complete dendritic filling and clear 3D visualization. Neurons with incomplete staining or visualization were excluded, and spine sampling focused on well-characterized spines orthogonal to the dendritic shaft. Data were analyzed using NeuroExplorer following (38).

### Protein measurement via Western Blot

Brain tissue from the chronic 3- and 6-month groups (each consisting of 10 mice, with approximately 50% female) was prepared for protein analysis. The prefrontal cortex (PFC) and whole hippocampus regions were sonicated in APL buffer from Qiagen AllPrep RNA/Protein extraction kit (Qiagen, Germany; Cat # 80404). Proteins were extracted from the brain homogenate. Protein concentrations were determined using the Bicinchoninic Acid (BCA) assay (Thermo Fisher Scientific, USA). Bovine serum albumin (BSA) standards were used to generate a standard curve, and absorbance was measured at 562 nm with a microplate reader.

Equal amounts of protein (20 μg) from each sample, as determined by the BCA assay, were separated on 10% SDS-PAGE stain-free gels (Bio-Rad Laboratories, #5678094). Following electrophoresis, the gels were imaged using the ChemiDoc system (Bio-Rad Laboratories, USA) to verify equal loading and quantify total protein quantification. Proteins were then transferred onto nitrocellulose membranes using a semi-dry transfer system. Membranes were blocked with 5% BSA in 1X TBST (Tris-buffered saline with 0.1% Tween 20) for 1 hour at room temperature. Membranes were incubated overnight at 4°C with primary antibodies targeting p-Tau or Tau (see **Supplementary Table 1** for details). After washing, membranes were incubated with HRP-conjugated secondary antibodies (Vector, #PI-1000 for rabbit and #PI-2000 for mouse) for 1 hour at room temperature. Signal detection was performed using an enhanced chemiluminescence (ECL) substrate (Thermo Fisher Scientific, USA) and the membrane was imaged in the ChemiDoc. Quantitative analysis of total protein and band intensity was conducted using ImageLab software.

### Study Design

Heterozygote PS19 mice and their WT littermate were studied at 3 months of age (corresponding to an early stage of phosphorylation of Tau; **Fig.1AB**) or at 6 months of age (corresponding to a more advanced stage of phosphorylation of Tau; **Fig.1CD**). At each age, the effect of acute or chronic (4 weeks) administration of GL-II-73 was investigated on working memory performances in the Y maze. In the chronic studies, brains were extracted for downstream analyses including morphology investigation using a Golgi staining technique, or western blot, 7 days after completion of the Y maze.

### Statistics

GraphPad Prism (v10.3.1) was employed for statistical analyses, and the results were graphically presented to convey mean ± SEM (**Supplementary Tables 2-4**). Morphological Golgi stain results were analyzed by Neurodigitech through two-way ANOVAs with Tukey’s multiple comparisons test. The Y-maze data underwent analysis via Kruskal-Wallis test, followed by uncorrected Dunnett’s multiple comparisons test. The p-tau/Tau ratio was calculated from the western blot relative expression of pTau and Tau, with the limitation that pTau levels in WT were not detectable. Therefore, relative expression was calculated using the HET group as a reference.

## RESULTS

### Chronic GL-II-73 treatment prevents neuronal shrinkage and reduced spine density in the PFC and CA1 of PS19 mice at 3 and 6 months of age

Morphological changes were quantified on pyramidal neurons located in the prefrontal cortex (PFC) and the CA1 of the hippocampus (HIP) using a Golgi-Cox staining technique on mice from the chronic studies at 3 and 6 months.

At 3 months of age, untreated HET mice showed a significant reduction in spine density in the PFC and CA1, corresponding to 80-81.5% of WT control levels (**Figure 2G**, top panel). This reduction was partially prevented by chronic administration (1 month) of GL-II-73 in both brain regions (**Supplementary Table 2**). The spine density of PS19 HET treated with GL-II-73 (HET+GL) were at 92.7% and 92.4% of WT levels in PFC and CA1 respectively, significantly higher than untreated HET mice, but still lower than WT mice. Similar reductions of spine density in untreated HET mice were observed at 6 months of age (79.3% and 69.4% of WT levels in PFC and CA1, respectively), which were in large part prevented by chronic GL-II-73 treatment (95.6% and 93.7% in PFC and CA1 of WT mice, respectively) (**Figure 2H; Supplementary Table 3**). The reduction in spine density in HET mice and partial reversal with GL-II-73 treatment was observed in apical and basal segments of the pyramidal neurons (**Supplementary** Figure 1) in both the PFC and CA1.

**Figure 2.**
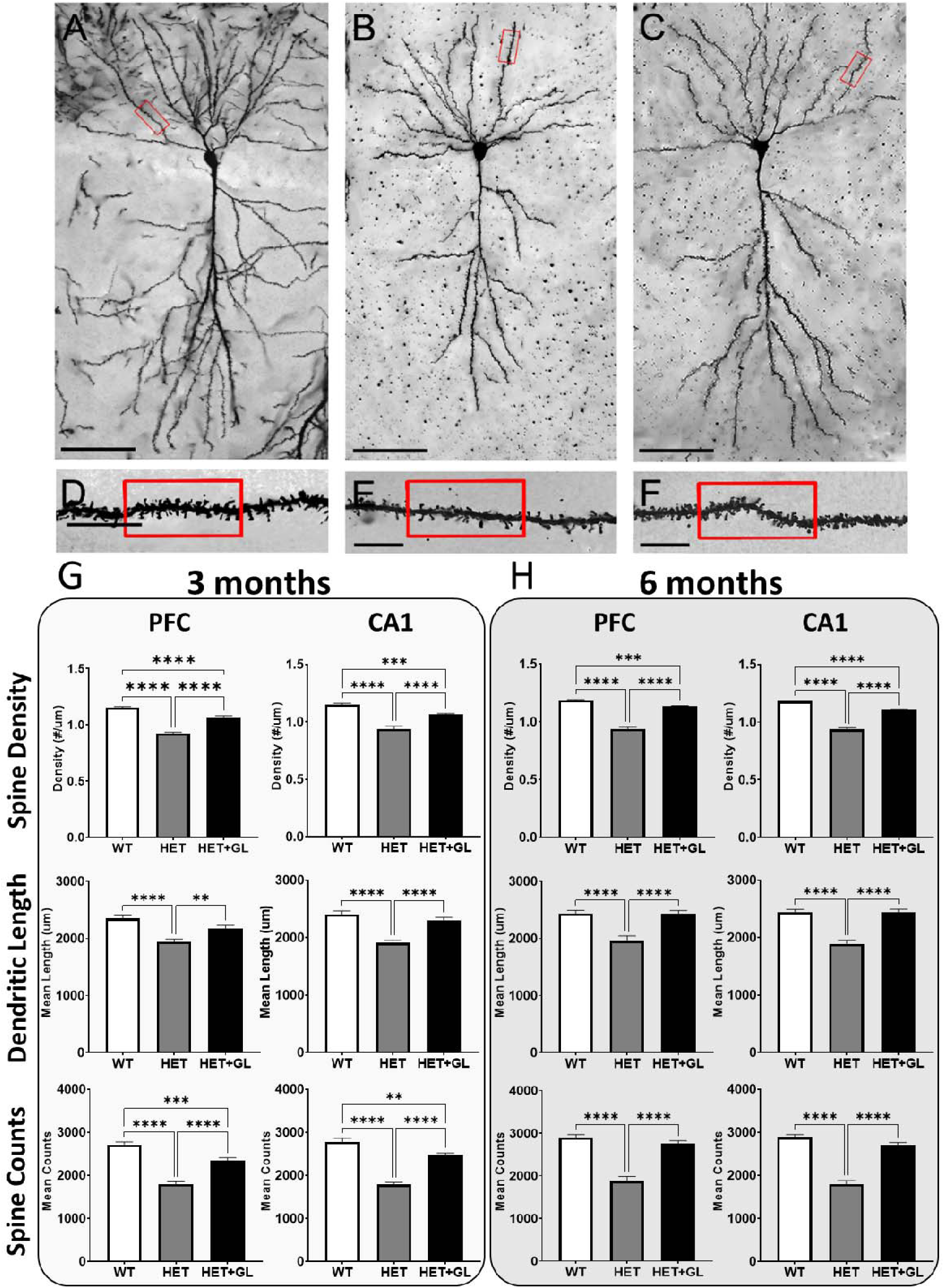
Chronic treatment with GL-II-73 reverses the reduced spine density, dendritic length and spine loss in the PFC and CA1 of PS19 mice at 3 and 6 months of age. Four brains per group were collected after chronic administration of GL-II-73 at 3 and 6 months. Six pyramidal neurons in the PFC (illustrated in **A-F**) and 6 principal cells of the CA1 of the HIP were isolated per animal and quantified for spine density, dendritic length and spine counts (PFC (**G**) and CA1 (**H**)). Untreated HET mice showed a significant reduction in spine density at 3 months in the PFC and CA1 which were partially reversed by chronic GL-II-73 (**G**). Similar effects were observed at 6 months (**H**). ***p<0.001, ****p<0.0001. Mean ±SEM.

Similar results were obtained for the analyses of dendritic length and spine counts (**Figure 1H** and **G**, middle and lower panels). Significant reductions of dendritic length were observed, corresponding to 82.6% and 79.3% of WT levels at 3 months and 80.2% and 77.5% of WT levels at 6 months in the PFC and CA1, respectively, which were reversed or prevented by GL- II-73 treatment. Dendritic length of mice treated with GL-II-73 were not significantly different from WT levels at 3 months (92.6% and 95.8% of WT levels in the PFC and CA1), and at 6 months (99.7% and 100% of WT levels in the PFC and CA1, respectively). Similarly, spine count reductions were observed in untreated HET mice at both age, and in both brain regions (66.2% and 64.3% of WT levels in the PFC and CA1 at 3 months, and 64.7% and 62.2% of WT levels in the PFC and CA1 at 6 months), which were prevented or reversed by GL-II-73 treatment (86.2% and 88.5% of WT levels in the PFC and CA1 at 3 months, and 95% and 93.6% of WT levels in the PFC and CA1 at 6 months).

### Chronic GL-II-73 does not affect the accumulation of phosphorylated Tau

The PFC and HIP from 9-10 animals that were not used for Golgi staining were collected to measure Tau and phospho-Tau levels after chronic administration of GL-II-73 at 3 and 6 months of age (**Supplementary** Figure 2**, and Supplementary Table 4**). There was no group differences in expression of Tau in the PFC or HIP at 3 and 6 months. However, there was a significant increase in pTau levels in HET mice and in GL-II-73-treated HET mice, compared to WT. Taking the native expression of Tau and the expression of pTau in the same samples, a ratio between pTau/Tau was calculated as an index of disease progression (**Figure 3**). HET mice exhibited a significant increase in pTau/Tau ratio (driven by increase in pTau level; **Supplementary** Figure 2) in the PFC and HIP at 3 months and at 6 months. Mice receiving chronic administration of GL-II-73 also exhibited higher ratio of pTau/Tau compared to WT, which were nominally lower in the 6-months of age group but not significantly different from untreated HET mice in both brain regions and at both ages.

**Figure 3.**
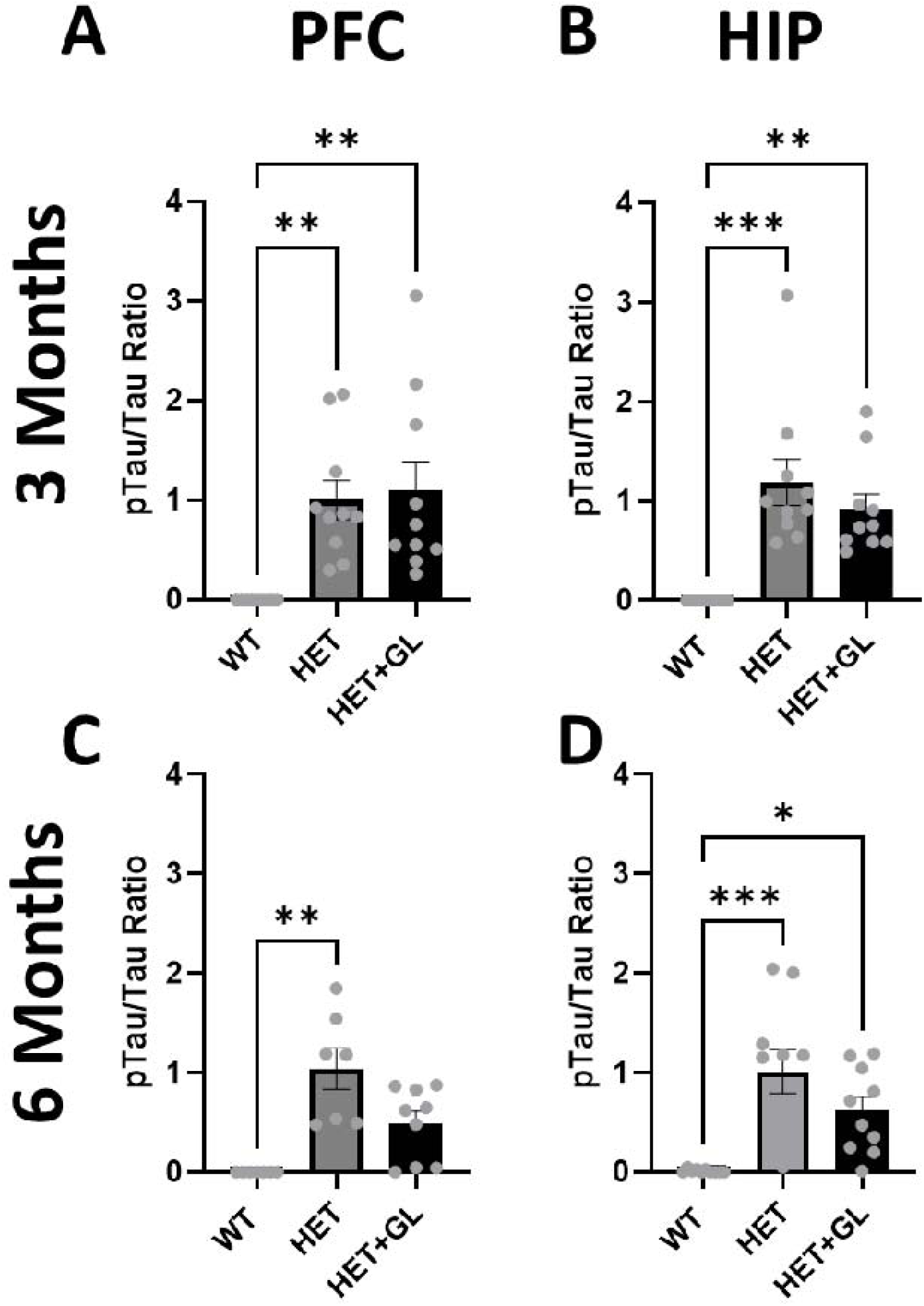
pTau/Tau ratio in the PFC and HIP from PS19 mice treated with GL-II-73 compared to untreated PS19 and WT controls. Using Western Blot targeting Tau and p-Tau, a p-Tau/Tau ration was calculated. HET mice at 3 months of age showed a higher ratio compared to WT (as expected) in the PFC (**A**) and HIP (**B**). HET mice receiving chronic GL-II-73 administration also showed higher ratio compared to WT, not different from untreated HET mice. At 6 months, the increased ratio in untreated HET compared to WT was confirmed in the PFC (**C**) and HIP (**D**). HET mice receiving GL-II-73 showed ratio not different from WT or HET in the PFC, and significantly higher than WT in the HIP. **p<0.01, ***p<0.001, ****p<0.0001. Mean ±SEM.

### Acute and chronic GL-II-73 treatment reverses working memory deficits

Mice were trained in the alternation task in the Y maze and tested at 3 months and 6 months, after either acute administration of GL-II-73, 30 minutes prior to testing, or after chronic administration in the drinking water for 4 weeks. In the groups of mice participating in the acute studies, PS19 HET mice showed a significant deficit in alternation compared to WT mice at 3 and 6 months (**Figure 4, and Supplementary Table 5**). Acute administration of GL-II-73 displayed a dose-dependent response with significant reversal of alternation deficits at 10 mg/kg and intermediate results at a lower 5 mg/k dose. Neither of the acute doses had significant effects at 6 months.

**Figure 4.**
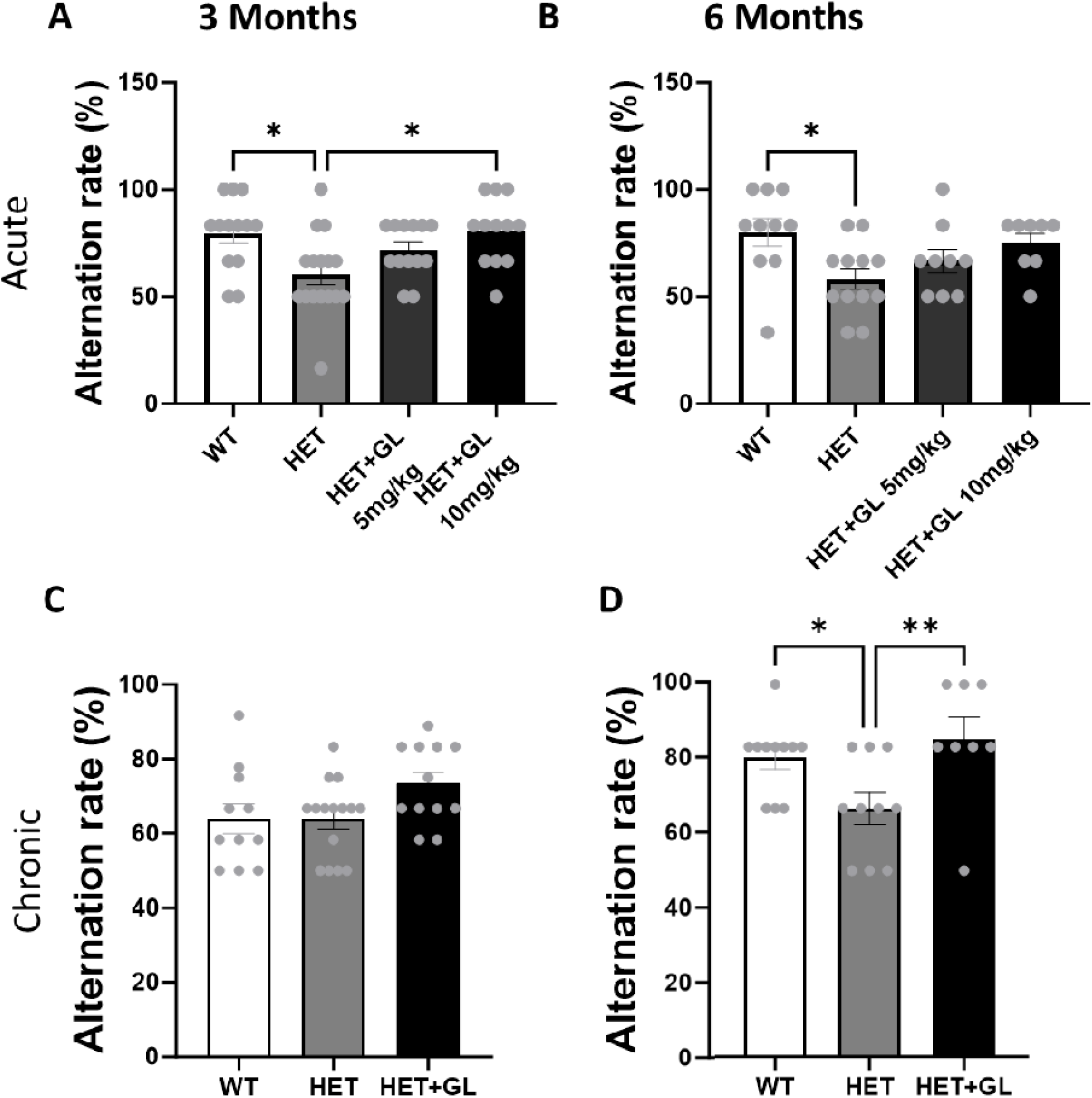
Effects of acute and chronic administration of GL-II-73 on alternation rate in PS19 HET mice. Alternation rate in the Y maze were measured after acute administration of GL-II-73 at 3 months (5 and 10 mg/kg dose) (**A**) and 6 months (10 mg/kg) (**B**). At 3 months, HET mice showed a reduction in alternation rate compared to WT. Acute treatment with 10mg/kg dose reversed this deficit, while 5 mg/kg did not significantly change alternation rate. At 6 months, untreated HET mice showed a significantly reduced alternation rate compared to WT. HET mice treated with GL-II-73 did not show significantly improved performances compared to untreated HET mice. Similarly, alternation rate in the Y maze were measured after chronic administration of GL-II-73 at 3 months (**C**) and 6 months (**D**). At 3 months, there was no difference between groups. At 6 months, untreated HET mice showed a significantly reduced alternation rate compared to WT. On the other hand, HET mice treated with GL-II-73 showed significantly higher alternation rate than untreated HET and comparable to WT levels. *p<0.05, **p<0.01. Mean ±SEM.

In the chronic administration studies (**Figure 5, and Supplementary Table 5**), alternation rates in the 3 months-old mice were not significantly different between groups. At 6 months, PS19 HET mice demonstrated a significant deficit in alternation rate deficit compared to WT, which was reversed by chronic administration of GL-II-73.

## DISCUSSION

In this study, we show that chronic administration of GL-II-73, a positive allosteric modulator of the α5-GABAAR subtype (*39*), prevents and reverses the loss of spine density in the prefrontal cortex and hippocampus of the PS19 mice in early and advanced stages of p-Tau accumulation. These effects occurred despite accumulation of p-Tau. In addition, improvement of cognitive functions and reversal of p-Tau-induced deficits, more specifically working memory, were observed, in particular at 3 months when given acutely, and 6 months when given chronically. Importantly, the beneficial effects on neuronal morphology and cognitive functions was independent of Tau-levels, since the chronic treatment did not significantly reduce the p-Tau accumulation.

These findings align with previous findings in other models, demonstrating that targeting the α5- GABAA-Rs with GL-II-73 has neurotrophic effects in the PFC and CA1, as well as cognitive benefits. Indeed, our group demonstrated that chronic administration of GL-II-73 prevents or reverses neuronal atrophy in animal models of chronic stress (*30*), aging (*31*), and amyloid deposition (*32*). Chronic treatment also provided relief of cognitive deficits in these models. This new finding in the context of p-Tau accumulation in mice further suggests that intervention with GL-II-73 in populations with tauopathies would provide neuronal support and would contribute to prevention or reversal of spine density loss and cognitive decline in conditions such as AD or FTD.

Importantly, the PS19 mouse model only expresses the 1N4R isoform, which represents 20-25% of the tauopathies in human, but is not the only isoform present in AD (*40*). A complete model of AD would require including the other isoforms reported to be present in AD (*40*), such as the various 3R isoforms. Interestingly, the MAPT 1N4R isoform expressed in the PS19 mice is present in other disorders with tauopathy, such as progressive supranuclear palsy (*41*), corticobasal degeneration (*42*), globular glial tauopathy (*43*) and FTD (*44*), which suggest they may be of relevance for intervention with α5-PAM for cognitive deficits and neurodegeneration in these disorders.

Targeting Tau pathologies has been of great interest over the years in the context of AD (*45*), as studies suggested that Tau shows a stronger correlation with symptom severity than does amyloid β (*46*). Interventions focusing on Tau protein modification failed in clinical trials over the years, like Salsalate (*47*), MR-8719, or recently LY3372689 that failed meeting primary endpoint in Phase 2, BIIB113 which was discontinued or ASN-51 Phase 2 trials halted in March 2025. Tau accumulation inhibitors are also investigated in clinical trials (LMTX also known as TRx0237) but their efficacy remains to be demonstrated. Immunotherapies are now the most prominent approaches used to target Tau pathologies in clinical trials (*8, 46, 48, 49*), with promising avenues (see (*45*) and (*46*) for reviews). Most trials are currently assessing safety and tolerability of such interventions. Overall, their impact on neuronal health or cognitive functions remains unclear. But since the Tau pathologies in AD is only one component of the disease, one can predict that targeting other aspects such as neuronal dysfunction can provide additional benefits than solely targeting Tau pathologies (*50*).

Targeting GABAergic neurotransmission, particularly through α5-GABAA R, offers a promising avenue in neurodegenerative disease research, such as AD and FTD. These receptors are predominantly expressed in hippocampal regions, brain areas critical for memory and cognition (*26*) and severely affected in AD. Dysregulation of GABAergic signaling in AD has been linked to cognitive decline, and modulating α5 GABAA-R activity could help restore inhibitory balance and improve cognitive function (*26*). Tauopathies were shown to generate hyperexcitability (*51*).

GL-II-73 may restore part of the excitation/inhibition balance by increasing GABAergic activity onto glutamatergic cells, thus stabilizing neuronal activity, altogether helping with cognitive functions and neuronal exhaustion and atrophy. Moreover, combining this approach with anti- Tau therapies could provide synergistic effects, as p-Tau accumulation contributes to neurodegeneration and cognitive impairments. By simultaneously addressing p-Tau accumulation, GABAergic dysfunction and neuronal loss, combination therapies have the potential to enhance treatment efficacy and slow disease progression.

Interestingly, the effects observed in this study were independent from p-Tau levels, as chronic treatment did not alter p-Tau levels in mice, neither at 3 months nor at 6 months. This observation can be due to the constitutive aspect of the model, where p-Tau is produced continuously in the PS19 mouse brain. However, despite this continuous accumulation of p-Tau, treatment with GL-II-73 was able to prevent spine density reduction and cognitive decline, suggesting strong efficacy despite the presence of the traditional hallmark of AD, similarly to what was previously demonstrated in a amyloid accumulation model (*32*).

This study has some limitations. As mentioned before, the model used only expresses the 1N4R isoform of Tau, while there are other isoforms present in AD and in other tauopathy. Therefore, confirming such effect in other models of tauopathy expressing the other isoforms, or a combination would further validate the efficacy of GL-II-73. However, in view of similar effects in models of stress (*29, 30*), aging (*31*), and amyloid deposition (*32*), the likelihood of generalization effects in other models is high. From a cognitive assessment standpoint, we only investigated the impact of working memory, when AD and other brain disorders associated with tauopathy are characterized by more cognitive impairments, such as episodic memory, executive function, attention and others. Investigating efficacy in these domains would represent an immediate next step to strengthen the potential of GL-II-73 as a pro-cognitive drug. In addition, the neurotrophic effect was observed in both regions investigated. Without a negative control, it is possible that the effect observed with the treatment is generalized throughout the brain, but similar findings were reported in the context of stress (*30*), aging (*31*), and amyloid deposition (*32*), suggesting that this effect on spine density is robust.

To conclude, we showed that chronic treatment with GL-II-73 demonstrated robust neurotrophic effects in the PS19 model of tauopathy in both early (2 months) and advanced (5 months) stages of p-Tau pathology, together supporting its clinical testing across stages of tauopathy, as stand- alone or combination therapy, and potentially prophylactically, to delay onset or slow down disease progression.

## FUNDING

This work has been funded by the Weston Brain Institute (TR192043).

## ACKNOWLEDGEMENTS

Authors thank Mehrab Ali for support with administrative tasks throughout the study, the CAMH animal facility staff for the caring for our animals over the study duration, the members of NeuroDigitech for their contribution to data generation, and all the animals that contributed to these studies.

## DISCLOSURES

JMC, MM, ES and TP are listed inventors on patents covering syntheses and use of the compound. ES is co-founder of Damona Pharmaceuticals, a biopharma dedicated to bring novel GABAergic compounds to the clinic. TP provides consulting services to Damona related to preclinical Research and Development. All other authors declare no conflicts of interest.

## Notes

### Summary of Updates

Update in text to develop the discussion points, and reorganized results for a better flow.

