## Supplementary for "α5-GABA-A Receptor Positive Allosteric Modulation prevents neuronal atrophy and cognitive decline independently of p-Tau accumulation in the PS19 mouse model"

**---Supplementary Material---**

Ravinder N. Dharavath^1+^, Ashley M. Bernardo^1+^, Cassandra Marceau-Linhares^1+^, Michael Marcotte^1^, Carla Mezo-Gonzalez^1^, Kayla Wong^1^, Celeste Pina-Leblanc^1^, Adrien Bouchet^1^, Dishary Sharmin^2^, Kamal P. Pandey^2^, James M. Cook^2^, Thomas D. Prevot^1,3,4*^, Etienne Sibille^1,3,4*^

^1^Campbell Family Mental Health Research Institute of CAMH, 250 college street, Toronto, ON, M5T 1R8 Canada

^2^Department of Chemistry and Biochemistry, University of Wisconsin–Milwaukee, 3210 N Cramer Street, 53211, WI, USA

^3^Department of Psychiatry, University of Toronto, 250 college street, Toronto, ON, M5T 1R8 Canada

^4^Department of Pharmacology and Toxicology, University of Toronto, Medical Sciences Building, 1 King's College Circle Room 4207, Toronto, ON, M5S 1A8, Canada

^+^These authors equally contributed to the study

*Corresponding Authors:

Etienne Sibille, Ph.D, CAMH, 250 College Street, room 134, Toronto, ON M5T 1R8, Canada

Thomas D. Prevot, Ph.D., CAMH, 250 College Street, room 131, Toronto, ON M5T 1R8, Canada

### **Supplementary Figures**

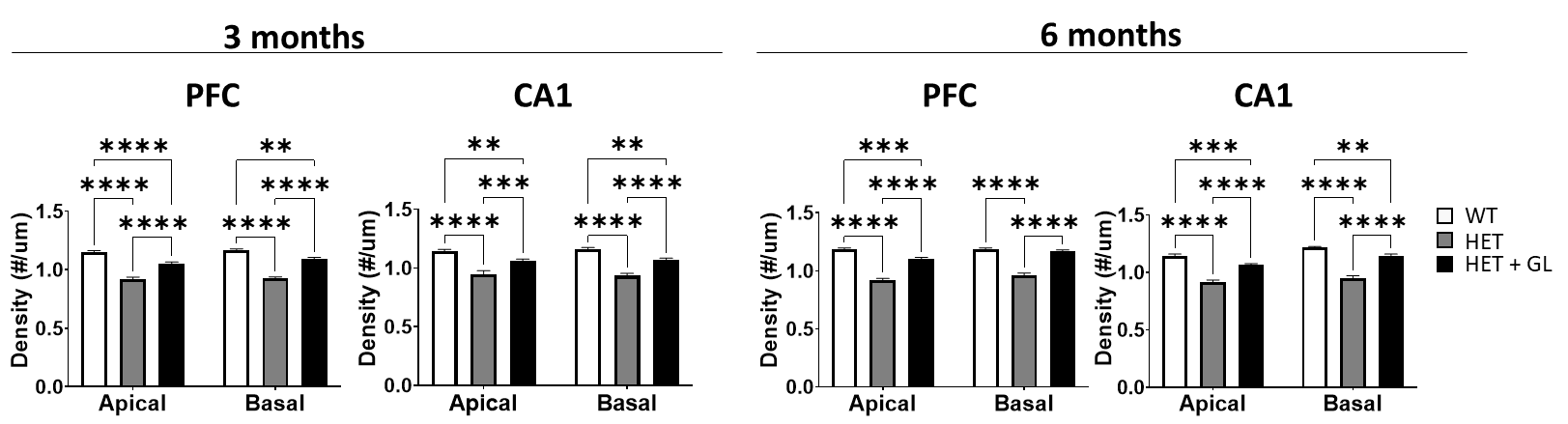

**Supplementary Figure 1.** Spine density in apical and basal segments

Four brains per group were collected after chronic administration of GL-II-73 at 3 and 6 months. Six pyramidal neurons in the PFC and 6 principal cells of the CA1 of the HIP were isolated per animal and quantified for spine density. Quantification of spine density was divided between the apical and the basal segment of the neuron. In both brain regions, at both ages and in both neuronal segments, untreated HET mice showed reduced spine density compared to WT. After chronic treatment with GL-II-73, HET+GL mice showed increased spine density compared the untreated HET. **p<0.01, ***p<0.001, ****p<0.0001. Mean ± SEM.

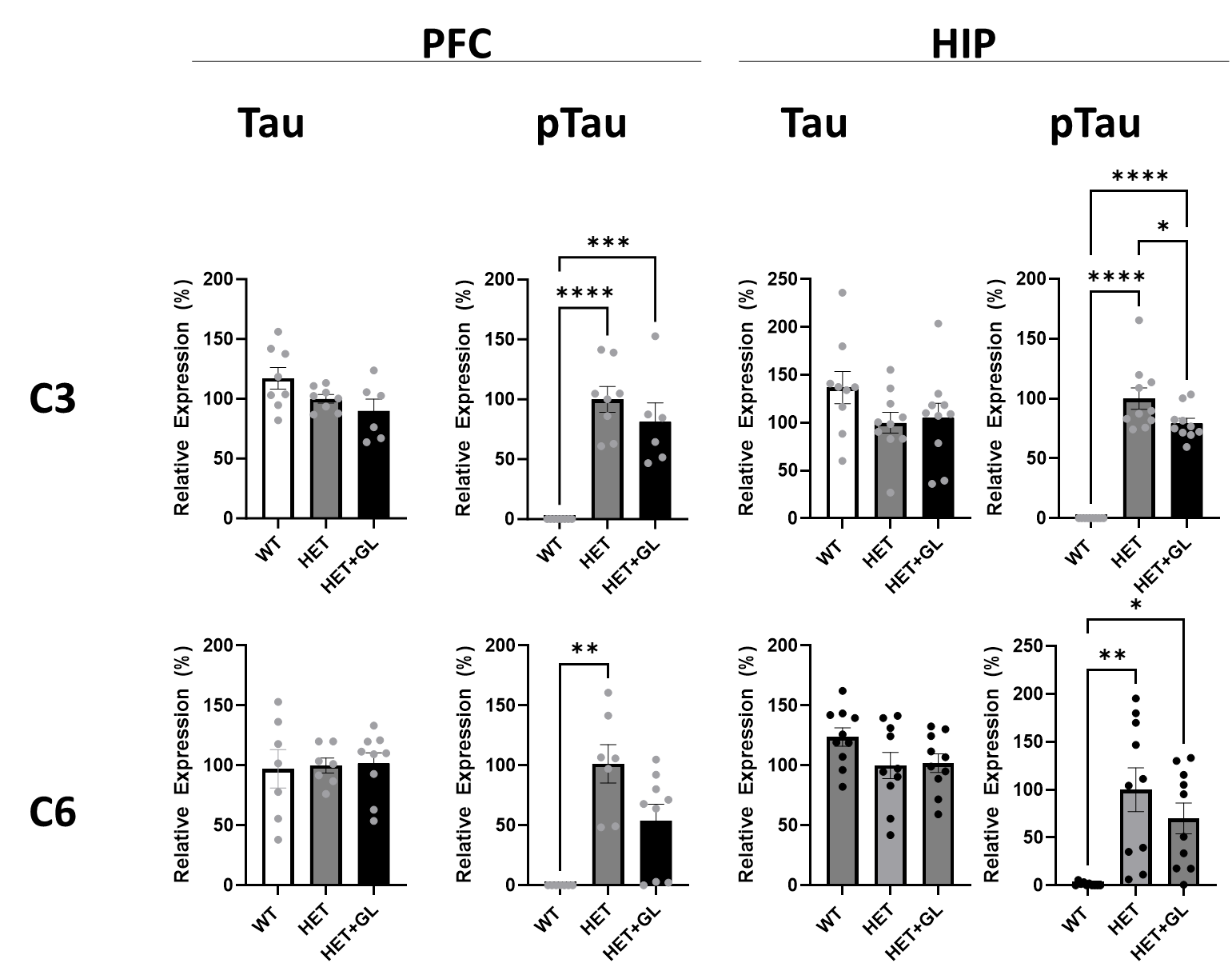

**Supplementary Figure 2.** Tau and pTau expression levels in the PFC and HIP of chronically treated PS19 HET mice in comparison to WT and untreated PS19 HET mice

Brain samples collected from 3 and 6 months old mice chronically treated with GL-II-73 or water were dissected to isolate the PFC and the hippocampus. Homogenates were purified to isolate proteins and western blot approaches were used to measure Tau levels, as well as phosphorylated Tau (pTau) levels. At both ages and in both brain regions, there was no difference found in the levels of Tau between groups. However, there was a significant increase in pTau levels in HET and HET+GL compared to WT at both ages and in both brain regions. In most cases, pTau levels in HET+GL mice were not different to the levels from HET mice. Only a significant reduction in HET+GL mice compared to untreated HET mice was found in the HIP of 3 months old treated mice. *p<0.05, **p<0.01, ***p<0.001. Mean ± SEM

### **Supplementary Tables**

**Supplementary Table 1**. Antibody List

| **Antibody** | **Manufacture & Cat #** | **Primary Dilution** | **Blocking Solution** |
| --- | --- | --- | --- |
| p-Tau- Thr181 (Rabbit) | Cell signaling | 1:1000 in 5%BSA | 5% BSA |
|  | #12885s |  |  |
| Tau (Mouse) | ProteinTech #23594303 | 1:2000 in 5% BSA | 5% Milk |

**Supplementary Table 2**. Statistical analyses of morphological changes in 3 months old mice

| **Age** | **Region** | **Population** | **Statistical Test** | **F-Value** | **p-value** |
| --- | --- | --- | --- | --- | --- |
| 3 Month | CA1 | Length (Basal) | Two-way ANOVA | F(2,70)=22.33 | **p<0.0001** |
|  |  |  | Tukey's test | " WT - Vehicle vs. PS19 - Vehicle" " WT - Vehicle vs. PS19 - GL" " PS19 - Vehicle vs. PS19 - GL" | **<0.0001** 0.7 **<0.0001** |
|  |  | Length (Apical) | Two-way ANOVA | F(1,140)=22.33 | p<0.001 |
|  |  |  | Tukey's test | " WT - Vehicle vs. PS19 - Vehicle" " WT - Vehicle vs. PS19 - GL" " PS19 - Vehicle vs. PS19 - GL" | **<0.0001** 0.7 **<0.0001** |
|  |  | Length (Total) | Two-way ANOVA | F(1,140)=22.33 | p<0.001 |
|  |  |  | Tukey's test | " WT - Vehicle vs. PS19 - Vehicle" " WT - Vehicle vs. PS19 - GL" " PS19 - Vehicle vs. PS19 - GL" | **<0.0001** 0.8 **<0.0001** |
|  |  | Spine count (Basal) | Two-way ANOVA | F (1, 210) = 6.855 | P = 0.0095 |
|  |  |  | Tukey's test | " WT - Vehicle vs. PS19 - Vehicle" " WT - Vehicle vs. PS19 - GL" " PS19 - Vehicle vs. PS19 - GL" | **<0.0001 0.0263 <0.0001** |
|  |  | Spine count (Apical) | Two-way ANOVA | F (1, 210) = 6.855 | P = 0.0095 |
|  |  |  | Tukey's test | " WT - Vehicle vs. PS19 - Vehicle" " WT - Vehicle vs. PS19 - GL" " PS19 - Vehicle vs. PS19 - GL" | **<0.0001 0.0015 <0.0001** |
|  |  | Spine count (Total) | Two-way ANOVA | F (2, 105) = 60.45 | **p<0.0001** |
|  |  |  | Tukey's test | " WT - Vehicle vs. PS19 - Vehicle" " WT - Vehicle vs. PS19 - GL" " PS19 - Vehicle vs. PS19 - GL" | **<0.0001 0.0021 <0.0001** |
|  |  | Density (Basal) | Two-way ANOVA | F (1, 210) = 0.05000 | P = 0.8233 |
|  |  |  | Tukey's test | " WT - Vehicle vs. PS19 - Vehicle" " WT - Vehicle vs. PS19 - GL" " PS19 - Vehicle vs. PS19 - GL" | **<0.0001 0.0017 <0.0001** |
|  |  | Density (Apical) | Two-way ANOVA | F (1, 210) = 0.05000 | P = 0.8233 |
|  |  |  | Tukey's test | " WT - Vehicle vs. PS19 - Vehicle" " WT - Vehicle vs. PS19 - GL" " PS19 - Vehicle vs. PS19 - GL" | **<0.0001 0.0062 <0.0001** |
|  |  | Density (Total) | Two-way ANOVA | F (2, 105) = 43.75 | **p<0.0001** |
|  |  |  | Tukey's test | " WT - Vehicle vs. PS19 - Vehicle" " WT - Vehicle vs. PS19 - GL" " PS19 - Vehicle vs. PS19 - GL" | **<0.0001 0.0006 <0.0001** |
|  | PFC | Length (Basal) | Two-way ANOVA | F (1, 210) = 3.685 | P = 0.0563 |
|  |  |  | Tukey's test | " WT - Vehicle vs. PS19 - Vehicle" " WT - Vehicle vs. PS19 - GL" " PS19 - Vehicle vs. PS19 - GL" | **0.0003** 0.3531 **0.0287** |
|  |  | Length (Apical) | Two-way ANOVA | F (1, 210) = 3.685 | P = 0.0563 |
|  |  |  | Tukey's test | " WT - Vehicle vs. PS19 - Vehicle" " WT - Vehicle vs. PS19 - GL" " PS19 - Vehicle vs. PS19 - GL" | **<0.0001 0.0099 0.003** |
|  |  | Length (Total) | Two-way ANOVA | F (2, 105) = 15.77 | P < 0.0001 |
|  |  |  | Tukey's test | " WT - Vehicle vs. PS19 - Vehicle" " WT - Vehicle vs. PS19 - GL" " PS19 - Vehicle vs. PS19 - GL" | **<0.0001** 0.0511 **0.0047** |
|  |  | Spine count (Basal) | Two-way ANOVA | F (1, 210) = 0.3664 | P = 0.5456 |
|  |  |  | Tukey's test | " WT - Vehicle vs. PS19 - Vehicle" " WT - Vehicle vs. PS19 - GL" " PS19 - Vehicle vs. PS19 - GL" | **<0.0001 0.0191 <0.0001** |
|  |  | Spine count (Apical) | Two-way ANOVA | F (1, 210) = 0.3664 | P = 0.5456 |
|  |  |  | Tukey's test | " WT - Vehicle vs. PS19 - Vehicle" " WT - Vehicle vs. PS19 - GL" " PS19 - Vehicle vs. PS19 - GL" | **<0.0001 <0.0001 <0.0001** |
|  |  | Spine count (Total) | Two-way ANOVA | F (2, 105) = 51.51 | P < 0.0001 |
|  |  |  | Tukey's test | " WT - Vehicle vs. PS19 - Vehicle" " WT - Vehicle vs. PS19 - GL" " PS19 - Vehicle vs. PS19 - GL" | **<0.0001 0.0002 <0.0001** |
|  |  | Density (Basal) | Two-way ANOVA | F (1, 210) = 2.000 | P = 0.1588 |
|  |  |  | Tukey's test | " WT - Vehicle vs. PS19 - Vehicle" " WT - Vehicle vs. PS19 - GL" " PS19 - Vehicle vs. PS19 - GL" | **<0.0001 0.013 <0.0001** |
|  |  | Density (Apical) | Two-way ANOVA | F (1, 210) = 2.000 | P = 0.1588 |
|  |  |  | Tukey's test | " WT - Vehicle vs. PS19 - Vehicle" " WT - Vehicle vs. PS19 - GL" " PS19 - Vehicle vs. PS19 - GL" | **<0.0001 0.0002 <0.0001** |
|  |  | Density (Total) | Two-way ANOVA | F (2, 105) = 148.4 | P < 0.0001 |
|  |  |  | Tukey's test | " WT - Vehicle vs. PS19 - Vehicle" " WT - Vehicle vs. PS19 - GL" " PS19 - Vehicle vs. PS19 - GL" | **<0.0001 <0.0001 <0.0001** |

**Supplementary Table 3**. Statistical analyses of morphological changes in 6 months old mice

| **Age** | **Region** | **Population** | **Statistical Test** | **F-Value** | **p-value** |
| --- | --- | --- | --- | --- | --- |
| 6 Month | CA1 | Length (Basal) | Two-way ANOVA | F (1, 210) = 2.712 | P = 0.1011 |
|  |  |  | Tukey's test | " WT - Vehicle vs. PS19 - Vehicle" " WT - Vehicle vs. PS19 - GL" " PS19 - Vehicle vs. PS19 - GL" | **<0.0001** 0.7594 **<0.0001** |
|  |  | Length (Apical) | Two-way ANOVA | F (1, 210) = 2.712 | P = 0.1011 |
|  |  |  | Tukey's test | " WT - Vehicle vs. PS19 - Vehicle" " WT - Vehicle vs. PS19 - GL" " PS19 - Vehicle vs. PS19 - GL" | **<0.0001** 0.7511 **<0.0001** |
|  |  | Length (Total) | Two-way ANOVA | F (2, 105) = 32.71 | P < 0.0001 |
|  |  |  | Tukey's test | " WT - Vehicle vs. PS19 - Vehicle" " WT - Vehicle vs. PS19 - GL" " PS19 - Vehicle vs. PS19 - GL" | **<0.0001** >09999 **<0.0001** |
|  |  | Spine count (Basal) | Two-way ANOVA | F (1, 210) = 1.065 | P = 0.3033 |
|  |  |  | Tukey's test | " WT - Vehicle vs. PS19 - Vehicle" " WT - Vehicle vs. PS19 - GL" " PS19 - Vehicle vs. PS19 - GL" | **<0.0001** 0.7105 **<0.0001** |
|  |  | Spine count (Apical) | Two-way ANOVA | F (1, 210) = 1.065 | P = 0.3033 |
|  |  |  | Tukey's test | " WT - Vehicle vs. PS19 - Vehicle" " WT - Vehicle vs. PS19 - GL" " PS19 - Vehicle vs. PS19 - GL" | **<0.0001 0.0396 <0.0001** |
|  |  | Spine count (Total) | Two-way ANOVA | F (2, 105) = 67.99 | P < 0.0001 |
|  |  |  | Tukey's test | " WT - Vehicle vs. PS19 - Vehicle" " WT - Vehicle vs. PS19 - GL" " PS19 - Vehicle vs. PS19 - GL" | **<0.0001** 0.1612 **<0.0001** |
|  |  | Density (Basal) | Two-way ANOVA | F (1, 210) = 24.07 | P < 0.0001 |
|  |  |  | Tukey's test | " WT - Vehicle vs. PS19 - Vehicle" " WT - Vehicle vs. PS19 - GL" " PS19 - Vehicle vs. PS19 - GL" | **<0.0001 0.0056 <0.0001** |
|  |  | Density (Apical) | Two-way ANOVA | F (1, 210) = 24.07 | P < 0.0001 |
|  |  |  | Tukey's test | " WT - Vehicle vs. PS19 - Vehicle" " WT - Vehicle vs. PS19 - GL" " PS19 - Vehicle vs. PS19 - GL" | **<0.0001 0.0002 <0.0001** |
|  |  | Density (Total) | Two-way ANOVA | F (2, 105) = 142.6 | P < 0.0001 |
|  |  |  | Tukey's test | " WT - Vehicle vs. PS19 - Vehicle" " WT - Vehicle vs. PS19 - GL" " PS19 - Vehicle vs. PS19 - GL" | **<0.0001 <0.0001 <0.0001** |
|  | PFC | Length (Basal) | Two-way ANOVA | F (1, 210) = 2.648 | P = 0.1052 |
|  |  |  | Tukey's test | " WT - Vehicle vs. PS19 - Vehicle" " WT - Vehicle vs. PS19 - GL" " PS19 - Vehicle vs. PS19 - GL" | **<0.0001** 0.931 **<0.0001** |
|  |  | Length (Apical) | Two-way ANOVA | F (1, 210) = 2.648 | P = 0.1052 |
|  |  |  | Tukey's test | " WT - Vehicle vs. PS19 - Vehicle" " WT - Vehicle vs. PS19 - GL" " PS19 - Vehicle vs. PS19 - GL" | **<0.0001** 0.8775 **<0.0001** |
|  |  | Length (Total) | Two-way ANOVA | F (2, 105) = 16.79 | P < 0.0001 |
|  |  |  | Tukey's test | " WT - Vehicle vs. PS19 - Vehicle" " WT - Vehicle vs. PS19 - GL" " PS19 - Vehicle vs. PS19 - GL" | **<0.0001** 0.9975 **<0.0001** |
|  |  | Spine count (Basal) | Two-way ANOVA | F (1, 210) = 0.005117 | P = 0.9430 |
|  |  |  | Tukey's test | " WT - Vehicle vs. PS19 - Vehicle" " WT - Vehicle vs. PS19 - GL" " PS19 - Vehicle vs. PS19 - GL" | **<0.0001** 0.9979 **<0.0001** |
|  |  | Spine count (Apical) | Two-way ANOVA | F (1, 210) = 0.005117 | P = 0.9430 |
|  |  |  | Tukey's test | " WT - Vehicle vs. PS19 - Vehicle" " WT - Vehicle vs. PS19 - GL" " PS19 - Vehicle vs. PS19 - GL" | **<0.0001** 0.0763 **<0.0001** |
|  |  | Spine count (Total) | Two-way ANOVA | F (2, 105) = 43.96 | P < 0.0001 |
|  |  |  | Tukey's test | " WT - Vehicle vs. PS19 - Vehicle" " WT - Vehicle vs. PS19 - GL" " PS19 - Vehicle vs. PS19 - GL" | **<0.0001** 0.4432 **<0.0001** |
|  |  | Density (Basal) | Two-way ANOVA | F (1, 210) = 12.00 | P = 0.0006 |
|  |  |  | Tukey's test | " WT - Vehicle vs. PS19 - Vehicle" " WT - Vehicle vs. PS19 - GL" " PS19 - Vehicle vs. PS19 - GL" | **<0.0001** 0.5777 **<0.0001** |
|  |  | Density (Apical) | Two-way ANOVA | F (1, 210) = 12.00 | P = 0.0006 |
|  |  |  | Tukey's test | " WT - Vehicle vs. PS19 - Vehicle" " WT - Vehicle vs. PS19 - GL" " PS19 - Vehicle vs. PS19 - GL" | **<0.0001 0.0003 <0.0001** |
|  |  | Density (Total) | Two-way ANOVA | F (2, 105) = 171.4 | P < 0.0001 |
|  |  |  | Tukey's test | " WT - Vehicle vs. PS19 - Vehicle" " WT - Vehicle vs. PS19 - GL" " PS19 - Vehicle vs. PS19 - GL" | **<0.0001 0.0008 <0.0001** |

**Supplementary Table 4.** Statistical analyses of the Tau, pTau and ratio levels in the PFC and HIP of mice at 3 and 6 months

| **Age** | **Region** | **Marker** | **Statistical Test** | **F-Value** | **p-value** |
| --- | --- | --- | --- | --- | --- |
| **3 Months** | **HIP** | **Tau** | Two-way ANOVA | F (2, 17) = 2.188 | P=0.1427 |
|  |  |  | Tukey’s | "WT vs. HET" "WT vs. HET+GL" "HET vs. HET+ GL" | **-**  **-**  **-** |
|  |  | **pTau** | Two-way ANOVA | F (2, 17) = 104.0 | **P<0.0001** |
|  |  |  | Tukey’s | "WT vs. HET" "WT vs. HET+GL" "HET vs. HET+ GL" | **<0.0001**; **<0.0001**; **0.0237** |
|  |  | **Ratio** | Two-way ANOVA | F (2, 17) = 16.62 | **P<0.0001** |
|  |  |  | Tukey’s | "WT vs. HET" "WT vs. HET+GL" "HET vs. HET+ GL" | **0.0001**; **0.0013**; 0.4235 |
|  | **PFC** | **Tau** | Kruskall Wallis | 19.9 | **<0.0001** |
|  |  |  | Dunn’s | "WT vs. HET" "WT vs. HET+GL" "HET vs. HET+ GL" | **0.0002**; **0.0005**; >0.9999 |
|  |  | **pTau** | Kruskall Wallis | 20.0 | **<0.0001** |
|  |  |  | Dunn’s | "WT vs. HET" "WT vs. HET+GL" "HET vs. HET+ GL" | **<0.0001**; **0.0019**; >0.9999 |
|  |  | **Ratio** | Kruskall Wallis | 19.9 | **<0.0001** |
|  |  |  | Dunn’s | "WT vs. HET" "WT vs. HET+GL" "HET vs. HET+ GL" | **0.0002**; **0.0005**; >0.9999 |
| **6 Months** | **HIP** | **Tau** | Two-way ANOVA | F (2, 18) = 2.196 | P=0.1401 |
|  |  |  | Tukey’s | "WT vs. HET" "WT vs. HET+GL" "HET vs. HET+ GL" | **-**  **-**  **-** |
|  |  | **pTau** | Two-way ANOVA | F (2, 18) = 9.179 | **P=0.0018** |
|  |  |  | Tukey’s | "WT vs. HET" "WT vs. HET+GL" "HET vs. HET+ GL" | **0.0016**; **0.0242**; 0.4296 |
|  |  | **Ratio** | Two-way ANOVA | F (2, 18) = 10.04 | **P=0.0012** |
|  |  |  | Tukey’s | "WT vs. HET" "WT vs. HET+GL" "HET vs. HET+ GL" | **0.0009**; **0.0364**; 0.2201 |
|  | **PFC** | **Tau** | Two-way ANOVA | F (2, 11) = 0.1899 | P=0.8297 |
|  |  |  | Tukey’s | "WT vs. HET" "WT vs. HET+GL" "HET vs. HET+ GL" | **-**  **-**  **-** |
|  |  | **pTau** | Two-way ANOVA | F (2, 11) = 8.757 | **P=0.0053** |
|  |  |  | Tukey’s | "WT vs. HET" "WT vs. HET+GL" "HET vs. HET+ GL" | **0.0040**; 0.0805; 0.0849 |
|  |  | **Ratio** | Two-way ANOVA | F (2, 11) = 7.687 | **P=0.0082** |
|  |  |  | Tukey’s | "WT vs. HET" "WT vs. HET+GL" "HET vs. HET+ GL" | **0.0066**; 0.1652; **0.0687** |

**Supplementary Table 5**. Statistical analyses of the Y maze data

| **Age** | **Dosing regimen** | **Statistical Test** | **Value** | **p-value** |
| --- | --- | --- | --- | --- |
| **3 Months** | **Acute** | **Kruskal Wallis** | 11.30 | **0.0102** |
|  |  |  | "WT vs. HET" "HET vs. HET+GL 5mg/kg" "HET vs. HET+ GL 10mg/kg" | **0.0149**; 0.3408; **0.0119** |
|  | **Chronic** | **Kruskal Wallis** | 4.456 | 0.1078 |
|  |  |  | "WT vs. HET" "WT vs. HET+GL" "HET vs. HET+ GL" | **-**  **-**  **-** |
| **6 Months** | **Acute** | **Kruskal Wallis** | 8.023 | **0.0455** |
|  |  |  | "WT vs. HET" "HET vs. HET+GL 5mg/kg" "HET vs. HET+ GL 10mg/kg" | **0.0329**; >0.9999; 0.1907 |
|  | **Chronic** | **Kruskal Wallis** | 8.187 | **0.0167** |
|  |  |  | "WT vs. HET" "WT vs. HET+GL" "HET vs. HET+ GL" | **0.0450**; 0.3538; **0.0059** |
